# Dendrite-soma interactions in cultured hippocampal neurons form non-random structural motifs with local presynaptic enrichment and strengthening

**DOI:** 10.64898/2026.05.15.725585

**Authors:** Yehonatan Greiner, Wolfgang Kurz, Melvin Dray, Gilad Lavi, Orly E. Weiss, Danny Baranes

## Abstract

Dendritic arbor morphology is shaped in part by interactions with neighboring dendrites, and its geometry strongly influences the spatial distribution and strength of synapses. These observations raise the possibility that local dendritic contacts help determine where synapses accumulate and strengthen. Previous work in cultured hippocampal neurons showed that dendrite–dendrite contact sites are non-random and associated with local synaptic clustering. Here we asked whether a different type of dendritic contact, formed between a dendrite and the soma of a neighboring neuron, behaves similarly. Using dissociated hippocampal cultures, immunofluorescence imaging, time-lapse microscopy, quantitative image analysis, stochastic spatial simulations, and minimal quantitative modeling, we identified three recurrent classes of dendrite-soma interactions (DSIs): dendrites crossing directly over a neighboring soma, growing tangentially along the soma perimeter, or contacting the proximal region where a neighboring dendrite emerges from the soma. These interactions were abundant, occurred exclusively between different neurons, and showed substantial structural persistence over several days. Their overall frequency exceeded stochastic predictions across culture densities, and two configurations - proximal and tangential contacts - were selectively enriched above random expectation, whereas soma-crossing contacts were largely consistent with stochastic overlap. DSI composition also changed over development, with proximal contacts becoming progressively more prevalent. At DSI sites, synaptophysin-positive puncta were significantly denser and more intense than on non-interacting dendritic segments, consistent with local enrichment and strengthening of presynaptic specializations. Minimal modeling further indicated that biased formation together with developmental stabilization explains the observed organization better than stochastic geometry alone. These findings identify DSIs as non-random structural motifs in cultured hippocampal networks and suggest that dendrite contact geometry can contribute to synaptic distribution and strengthening.

## Introduction

Dendritic arbors are now understood not as passive receivers of synaptic input, but as active and compartmentalized computational structures that shape how neuronal signals are sampled, integrated, and transformed into output [1–3]. The position of synaptic inputs along the dendritic tree is therefore functionally consequential rather than anatomically incidental [4,5], because local dendritic properties and branch-specific organization strongly influence the integration of nearby inputs [6]. In hippocampal pyramidal neurons, synaptic organization is distributed non-uniformly across somatic, proximal, and distal compartments. These distinct dendritic domains participate in spatially structured input organization rather than uniform synaptic sampling [7,8]. Dendritic architecture and synaptic distribution should therefore be considered together as integral components of the spatial logic of neuronal connectivity.

Beyond the absolute number of synapses, their spatial arrangement along dendrites is itself an important organizational variable [9,10]. Synaptic inputs are often organized in local clusters or functional dendritic domains, and such clustering has been proposed to enhance local cooperativity, promote nonlinear dendritic integration, and increase the computational capacity of individual branches [9,10]. Recent *in vivo* work further showed that clustered synapses emerge during development within confined dendritic segments and that co-active neighboring inputs are associated with synaptic stabilization and potentiation, linking spatial proximity to local strengthening of synaptic connections [4]. At the same time, plasticity along dendritic segments is not simply additive: neighboring synapses can interact cooperatively or competitively over local spatial scales, indicating that spatial organization can influence which inputs are selectively strengthened and retained [11,12]. Synaptic clustering should therefore be viewed not only as a descriptive pattern of input placement, but as a plausible mechanism linking dendritic spatial organization to local synaptic specialization and plasticity.

Dendritic patterning is shaped by a combination of intrinsic programs and extrinsic cues, including transcriptional and cytoskeletal regulation, adhesion-mediated interactions, and local synaptic or network activity, rather than by unconstrained branch growth alone [13,14]. Environmental constraints in this context also include the spacing and overlap of neighboring dendritic fields, because arbors developing within shared territory must balance coverage with avoidance of redundant innervation, a relationship classically described in tiling and arbor-spacing studies [15]. At the same time, self-avoidance mechanisms indicate that dendrites do not respond uniformly to all nearby branches: recent work on γ-protocadherins showed that developing dendrites selectively recognize and avoid self-contacts rather than indiscriminately repelling all neighboring processes [14]. These frameworks explain important aspects of single-arbor development, but they leave open a different level of organization that becomes apparent when dendrites enter sustained spatial relationships with neighboring neuronal compartments, where local contact architecture may itself contribute to synaptic patterning.

In cultured hippocampal networks, local neurite appositions can be visualized and quantified over developmental time with an optical and experimental access that is difficult to achieve in intact tissue, making primary cultures a useful system for testing whether recurring contact motifs reflect simple crowding or spatially biased wiring rules. Within this experimentally accessible framework, previous work in cultured hippocampal networks showed that recurrent dendrite–dendrite contact sites are predominantly non-self configurations, formed between processes of different neurons rather than within single arbors, and that their incidence can exceed stochastic expectations derived from random dendritic arrangements [16,17]. These findings suggest that local dendritic contacts in culture may represent regulated spatial motifs rather than incidental overlaps generated by network density alone. Importantly, the same studies further showed that such contact sites are associated with synaptic clusters whose density is higher than that of synapses located on non-intersecting dendritic segments, and that synapses positioned near these intersections exhibit greater presynaptic release efficacy than synapses outside intersection sites [16–18]. Together, these observations indicate that non-self dendrite–dendrite contacts can organize both the local accumulation and the local functional strengthening of synapses [16–18]. However, dendrites in neural tissue and in culture can also contact neuronal compartments other than other dendrites, raising the question of whether dendrite–cell body contacts may occur as well and whether they share organizational and functional properties with dendrite–dendrite interactions.

Dendrite–cell body contacts represent a distinct class of inter-neuronal apposition, because the soma defines a different geometric and compartmental context than a dendritic branch and could therefore impose different structural or synaptic constraints on a contacting dendrite. In some systems, such contacts are already known to occur and to carry functional significance [19]. In mouse retina, for example, AII amacrine cell dendrites form selective dendro-somatic synapses onto OFFα ganglion cells, showing that dendrite-soma contacts can support highly specific and physiologically meaningful connectivity patterns [19]. Developmental studies further indicate that dendrite-soma contacts can participate in tissue patterning itself: neighboring starburst amacrine cell dendrites transiently contact homotypic somata and contribute to mosaic spacing by excluding somata from settling within an adjacent dendritic territory [20]. Together, these findings suggest that dendrite–cell body contacts may define spatial microdomains in which local geometry biases the accumulation, stabilization, and strengthening of synaptic inputs, consistent with broader views that clustered synaptic organization on dendrites contributes to local integration and plasticity [5,10]. This, in turn, raises the question of whether comparable heteroneuronal dendrite-cell body appositions arise in hippocampal cultures as recurrent spatial motifs, whether they are better explained by random overlap or by regulated local wiring, and whether they may also represent sites of local synaptic enrichment, as observed at dendrite–dendrite intersections [16–18].

Against this background, we asked whether heteroneuronal dendrite–cell body contacts that arise in cultured hippocampal networks constitute controlled and regulated motifs rather than incidental overlaps generated by dense growth. We therefore sought to identify and characterize these interactions, here termed dendrite–soma interactions (DSIs), to determine whether they form in a non-random manner, and whether synapses become locally enriched at such sites. We found that DSIs are abundant, structurally diverse, and non-randomly organized, and that their prevalence changes over development. Moreover, representative DSI sites were associated with local presynaptic enrichment and increased synaptic strength, supporting the conclusion that dendrite–cell body contacts constitute a regulated spatial motif that may contribute to the local organization of synaptic connectivity and extend current views of how dendritic contact geometry shapes neural wiring.

## Materials and Methods

### Culture Substrate Preparation and Coating

To modulate the adhesion strength of hippocampal cells to the substrate, 12 mm glass (41001112, Karl Hecht Assistent) coverslips were modified with distinct coating materials to provide varied growth environments: Poly-D-Lysine (PDL, P7280, Sigma-Aldrich) was dissolved in Hank’s Balanced Salt Solution (H6648, Sigma-Aldrich) to a final concentration of 0.02 mg/ml solution. 100 µl were applied onto each coverslip and incubated overnight at 4° C. Before culturing, PDL-coated coverslips were washed once with double-distilled water (DDW) and air-dried in the biological hood. Another set of glass coverslips was coated with 0.085 µg/ml Collagen Type I (08-115, Merck Millipore) dissolved in 1X Phosphate buffer saline (1X PBS) and incubated at room temperature (RT) for one hour. Next, the solution residues were discarded and the coverslips were allowed to air-dry. For calcium carbonate coating, CaCO_3_ powder (GK3384, Glentham) was suspended in DDW at a concentration of 5 mg/ml. 100 µl of the mixture were applied to the coverslips, which were then heated on a heating plate to 80°C, until water evaporation, resulting in the formation of a uniform CaCO_3_ particle layer on the coverslip surface. The CaCO_3_-coated coverslips were then autoclaved. Following sterilization, CaCO_3_-coated coverslips were subsequently coated with PDL according to the protocol described above.

### Hippocampal Cell Cultures

Primary hippocampal cultures were prepared from Sprague Dawley rat pups, 0-3 days, in accordance with previously established protocols [21]. The hippocampi were dissected, enzymatically treated with 0.25% trypsin (15-090-046, Gibco) for 30 min at 37 °C, followed by mechanical trituration. Cells were plated at a density of 6 × 10⁵ cells/ml onto 12 mm glass coverslips pre-coated as described above. Cultures were initially maintained in Minimum Essential Medium (M2279, Sigma-Aldrich) supplemented with 10% heat-inactivated fetal bovine serum (Sigma-Aldrich), 1% L-glutamine (G7513s, Sigma-Aldrich), and 0.8% D-glucose (G8769, Sigma-Aldrich). The following day, the medium was replaced with serum-free medium containing 45% Minimum Essential Medium, 40% Dulbecco’s Modified Eagle Medium (D5796, Sigma-Aldrich), 10% F12 Ham nutrient mixture (N4888, Sigma-Aldrich), 0.25% (w/v) BSA (A4503, Sigma-Aldrich), 0.75% D-glucose (G8769, Sigma-Aldrich), 0.5% B27 supplement (17504-044, Gibco), 0.25% L-glutamine (G7513, Sigma-Aldrich), 0.01% kynurenic acid (K3375, Sigma-Aldrich), and 0.01% of a mixture containing 70% uridine (U0020, TCI) and 30% fluoro-deoxy-uridine (D2235, TCI). The cultures were maintained at 37 °C in a humidified incubator with 10% CO_2_ for the indicated time points, depending on the experimental design.

### Immunofluorescence

Cells were fixed with 4% paraformaldehyde (P6148, Sigma-Aldrich) for 10 min at RT, and washed four times with 1X PBS. The cells were then permeabilized with 0.25% Triton X-100 (H5142, Promega) for 5 min at RT, and blocked for one hour in 3% normal goat serum containing 0.1% Triton X-100, at RT. Blocking was followed by incubation of the cells (overnight, 4 °C) with the following primary antibodies: Rabbit anti-Neurofilament M (NFM, 0.002 mg/ml, AB1987, Merck Millipore), Mouse anti-microtubule-associated protein 2 (MAP2, 0.0025 mg/ml, AB5622, Merck Millipore), and Rabbit anti-synaptophysin (0.001 mg/ml, ab32127, Abcam). Cells were washed three times with 1X PBS, followed by incubation (one hour, RT) with the secondary antibodies Alexa Fluor 555 conjugated-Goat anti-rabbit IgG (0.004 mg/ml, A21429, Invitrogen), and Alexa Fluor 488 conjugated-Goat anti-mouse IgG (0.004 mg/ml, A11001, Invitrogen). Samples were then washed five times with 1X PBS, and mounted onto microscopic slides using Fluoromount (F4680, Sigma-Aldrich), supplemented with 2.5% 1,4-Diazabicyclo[2.2.2]octane (D27802, Sigma-Aldrich) as an antifade agent.

### Fluorescence Microscopy

Immunofluorescently stained samples were imaged using a Zeiss Axio Observer Z1 microscope equipped with 10x/0.30 Plan-Neofluar and 40x/1.30 Plan-Apochromat oil immersion objectives. Images were acquired with an AxioCam MRm CCD camera controlled by ZEN 2012 SP5 software. For the stability assay, an Olympus IX81 live-cell imaging system equipped with a PRIME BSI Express camera and UPlanFL N objectives (10x/0.30 Ph1 and 20x/0.50 Ph1) was used. The system was maintained at 37 °C with 5% CO_2_ in a humidified environmental chamber. Daily phase-contrast images of predefined fields were captured using a motorized stage and cellSens Dimension software.

### Confocal Microscopy and 3D Reconstruction

For 3D visualization of DSIs, Z-stacks of MAP2-labeled cells were captured using a Zeiss Axio Observer Z1 confocal system (40x/1.30 oil immersion objective). Excitation was performed with a 488 nm laser, and fluorescence was detected using PMTs. Acquisition parameters included 2x line averaging, a 1 Airy Unit (AU) pinhole, and a voxel size of 0.31×0.31×0.50 µm. Post-acquisition, raw stacks underwent deconvolution (Richardson–Lucy algorithm, 15 iterations) via the DeconvolutionLab2 plugin in Fiji using a theoretical point spread function. Final 3D reconstructions were produced using the Fiji 3D Viewer plugin with standardized threshold settings.

### Morphological Identification and Classification of DSIs

To morphologically identify and classify DSIs, MAP2-labeled cells were used to visualize neuronal architecture. To resolve overlapping dendrites and somatic boundaries, raw images were processed using the Contrast Limited Adaptive Histogram Equalization (CLAHE) algorithm in Fiji (ImageJ v2.16.0). Somas in dense aggregates or those with ambiguous boundaries were excluded from the analysis. For pyramidal neurons, DSI interactions were further sub-classified based on their localization to apical or basal dendritic segments. Identified DSIs were then counted manually.

### In Silico Modeling of Random Dendritic Distributions

To determine whether the observed frequency of DSIs exceeded what would be expected by chance, a custom graphical user interface (SWS-GUI) was developed in MATLAB (R2024a) to simulate simplified neuronal networks and quantify dendrite–soma interactions. Neuronal somas were modeled as non-overlapping circles randomly distributed within a rectangular working field. Dendritic processes were generated as cubic Bézier curves with randomized parameters governing length, orientation, and branching. Spatial scaling was calibrated such that one soma diameter corresponded to 10 µm. Dendrite–soma interactions in the simulation were classified using the same criteria established for the biological cultures. All simulations were generated using deterministic pseudorandom routines to ensure reproducibility across runs.

### Synaptic Quantification

Quantitative analysis of synaptic distribution and synaptophysin levels was performed by defining circular regions of interest (ROI, 5 µm diameter) over dendritic intersections, proximal dendritic segments, and intersegment shafts. Presynaptic puncta were identified using the Find Maxima algorithm in Fiji (ImageJ v2.16.0) with a fixed noise tolerance applied uniformly across all conditions. The Mean Gray Value was calculated exclusively within puncta-specific masks. To capture local variance, statistical analysis was conducted directly at the Individual ROI level, utilizing a one-way ANOVA.

### Pharmacological Blockade of Synaptic Activity

To assess whether synaptic activity influences DSI frequency, activity was pharmacologically modulated by supplementing the culture medium with either the NMDA receptor antagonist DL-2-amino-5-phosphonopentanoic acid (APV, 50 µM) or the AMPA/kainate receptor antagonist 6-cyano-7-nitroquinoxaline-2,3-dione (CNQX, 10 µM). These treatments were maintained for 48 hours prior to imaging and analysis.

### Statistical Analysis

For DSI stability, frequency per dendrites, and synaptic analyses, data reflect independent biological replicates. For other parameters, to account for local variance across large population sizes (N = 27–228), samples were pooled and randomly assigned to three representative subgroups for analysis. Statistical testing was performed using GraphPad Prism (v10.0). Normality was assessed, and comparisons were made using unpaired two-tailed t-tests (Welch’s correction) or one-way ANOVA with Tukey’s HSD test. Statistical significance was set at p < 0.05, with results presented as mean ± SEM.

### Minimal quantitative modeling of DSI structural organization and synaptic enrichment

To interpret the structural and synaptic datasets, we applied minimal quantitative models to test whether the observed DSI subtype distribution and synaptic puncta enrichment could be explained by stochastic geometry alone.

For structural organization, the observed frequency of subtype *i* at density *d* and developmental stage *t* was modeled as:

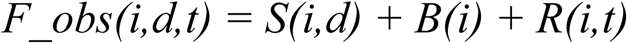

where *S(i,d)* is the stochastic expectation derived from the random spatial simulation, *B(i)* is a biased-formation term, and *R(i,t)* is a developmental refinement term. Based on this formulation, four nested models were compared:

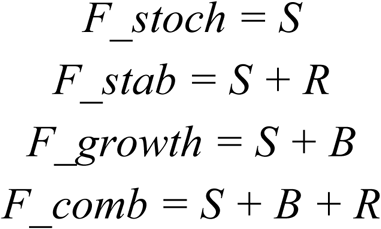

Model performance was compared using residual sum of squares (RSS):

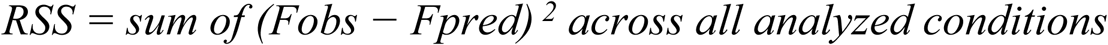

For synaptic enrichment, puncta abundance at DSI sites was modeled as:

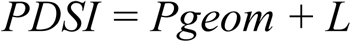

where *P_geom* is the puncta count expected from simple geometric overlap and L is an additional local contribution.

Assuming similar baseline puncta density in the interacting compartments, the geometric expectation was approximated as:

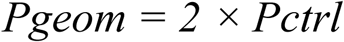

where *Pctrl* is the puncta count measured at a non-interacting control dendritic segment. The normalized enrichment ratio was defined as:

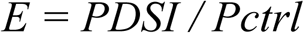

For this ratio, *E* = 2 represents the value expected from simple geometric superposition alone.

All modeling analyses were used as minimal descriptive comparisons of the quantified experimental data.

## Results

The experiments described below were designed to identify and quantify the occurrence of DSIs, to characterize their behavior under various manipulative conditions, and to examine their association with synaptic connection clustering. The study was conducted using immunofluorescence labeling of hippocampal neurons grown in culture for 7 to 21 days, followed by high-resolution imaging and quantitative image analysis.

### DSIs are morphologically variable, abundant and stable self-avoiding structural motifs

To define the structural nature of DSIs, we performed immunofluorescence staining for microtubule-associated protein 2 (MAP2) in hippocampal cultures (1-3 weeks old) Figure 1 establishes the morphological framework of DSIs, demonstrating that these contacts occur in distinct structural configurations between dendrites and neighboring somas across the cultured network.

**Fig. 1.**
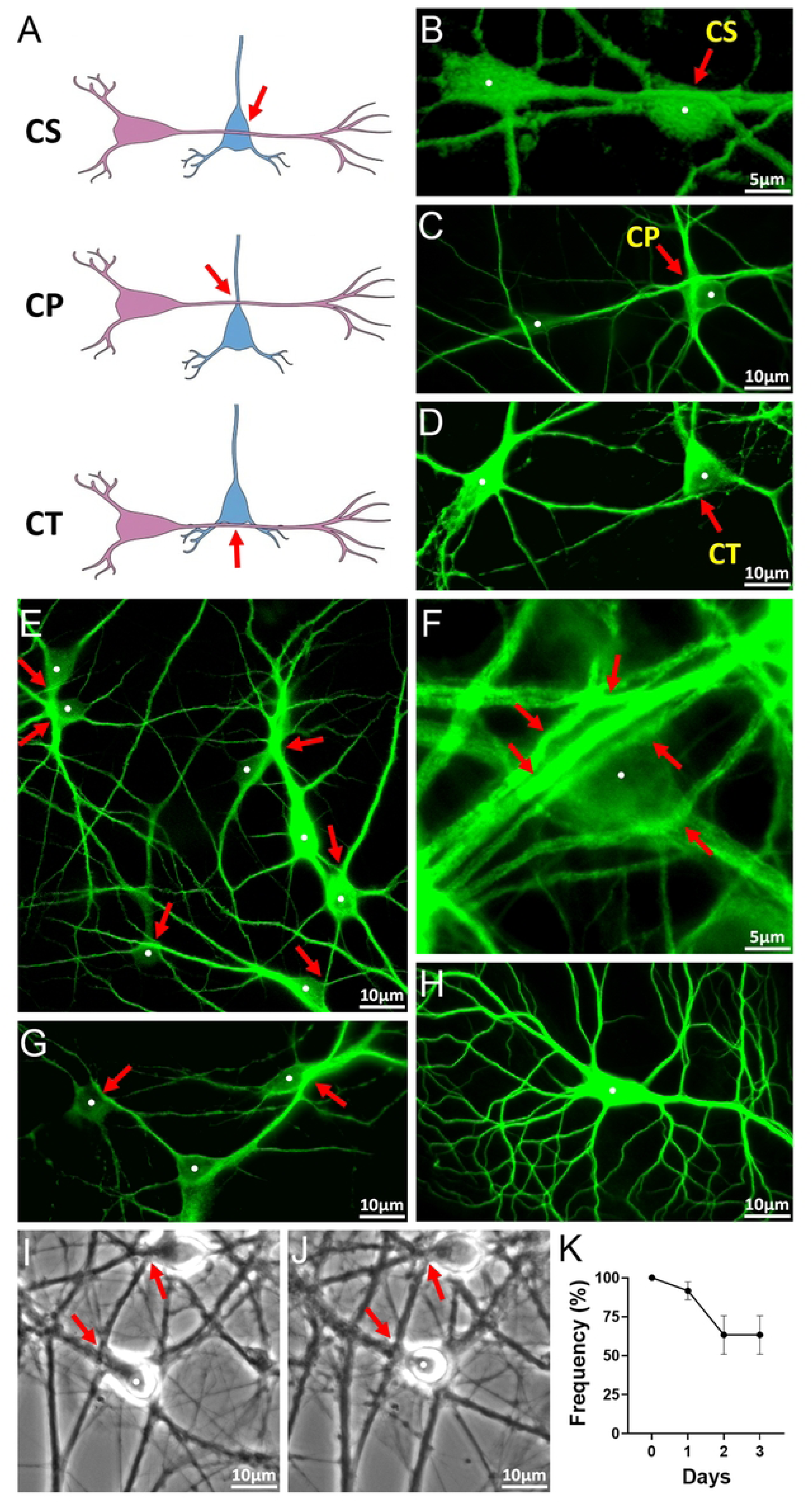
Morphological motifs and structural stability of dendrite–soma interactions (DSIs). Show hippocampal cultures at 7 DIV immunostained for MAP2 to visualize dendrites and cell bodies. Red arrows indicate sites of DSIs; white dots indicate cell bodies. (A–D) Representative schematics and immunofluorescence images illustrating three DSI morphological categories: Crossing-Soma (CS; A, B), Crossing-Proximal (CP; A, C), and Crossing-Tangential (CT; A, D). (E) Representative region of hippocampal culture showing frequent DSIs. (F) Example of a single soma hosting multiple DSIs. (G) An individual neuron forming multiple DSIs through different dendritic branches. (H) Absence of interactions between a neuron’s own dendrites and its soma. (I, J) Phase-contrast time-lapse imaging showing persistence of DSI sites from Day 0 (I) to Day 3 (J). (K) Quantification of DSI persistence over time, showing an initial decline followed by stabilization which remained stable through Day 3.

We identified three distinct morphological categories of DSIs, termed Crossing-Soma (CS), Crossing-Proximal (CP), and Crossing-Tangential (CT) (Figures 1A–D). In CS motifs, a dendrite crossed directly over the neighboring soma (Figures 1A and 1B). CP interactions were characterized by a dendrite crossing the proximal segment of a neighboring dendrite at its point of origin from the soma (Figures 1A and 1C). CT motifs involved a tangential crossing where the dendrite contacted the soma at a location distinct from any dendritic exit point, at the periphery of the soma (Figures 1A and 1D).

At the population level, these interactions were frequently observed across the hippocampal culture. For example, a representative area in culture shows DSIs occurring in high frequency (6 DSIs /7 somas, Figure 1E). Moreover, somas were found to host multiple DSIs from various neighboring dendrites (Figure 1F). Furthermore, individual neurons often established multiple DSIs through different dendritic branches (Figure 1G). Notably, neurons did not exhibit interactions between their own dendrites and their own somas (Figure 1H), indicating that DSIs occur exclusively between dendrites and the somas of neighboring neurons.

To evaluate structural stability, the persistence of DSI sites was monitored longitudinally over a three-day period across 10 individual neurons (Figures 1I and 1J). For each daily time point, the percentage of surviving DSIs was calculated relative to Day 0. Analysis revealed a gradual decline in interaction frequency during the first 48 hours, which stabilized at 62.33% by Day 2 and remained unchanged through Day 3 (Figure 1K). These values represent the mean survival rate of DSIs across the observed population over the three-day period.

The observed structural stability and abundance of DSIs prompted us to examine whether their occurrence reflects non-random organization within the neuronal network. To address this, we compared the experimental frequency of DSIs with predictions from a stochastic spatial simulation of cultured neuronal networks.

### DSIs occur more frequently than predicted by stochastic spatial simulations of cultured neuronal networks

To determine whether DSIs result from biological organization or from random interactions, we compared DSI occurrence in 7 day old cultures to that generated by a custom Soma Network Simulation (SWS-GUI) of randomized neuronal networks. The SWS-GUI modeled somas circles and dendrites as cubic Bézier curves within a defined simulation field (Figure 2A; see full description in the Methods section). Importantly, simulated interactions were classified into CS, CP, and CT categories using the same morphological criteria established for the biological cultures, ensuring a direct comparison between experimental and stochastic datasets.

**Fig. 2.**
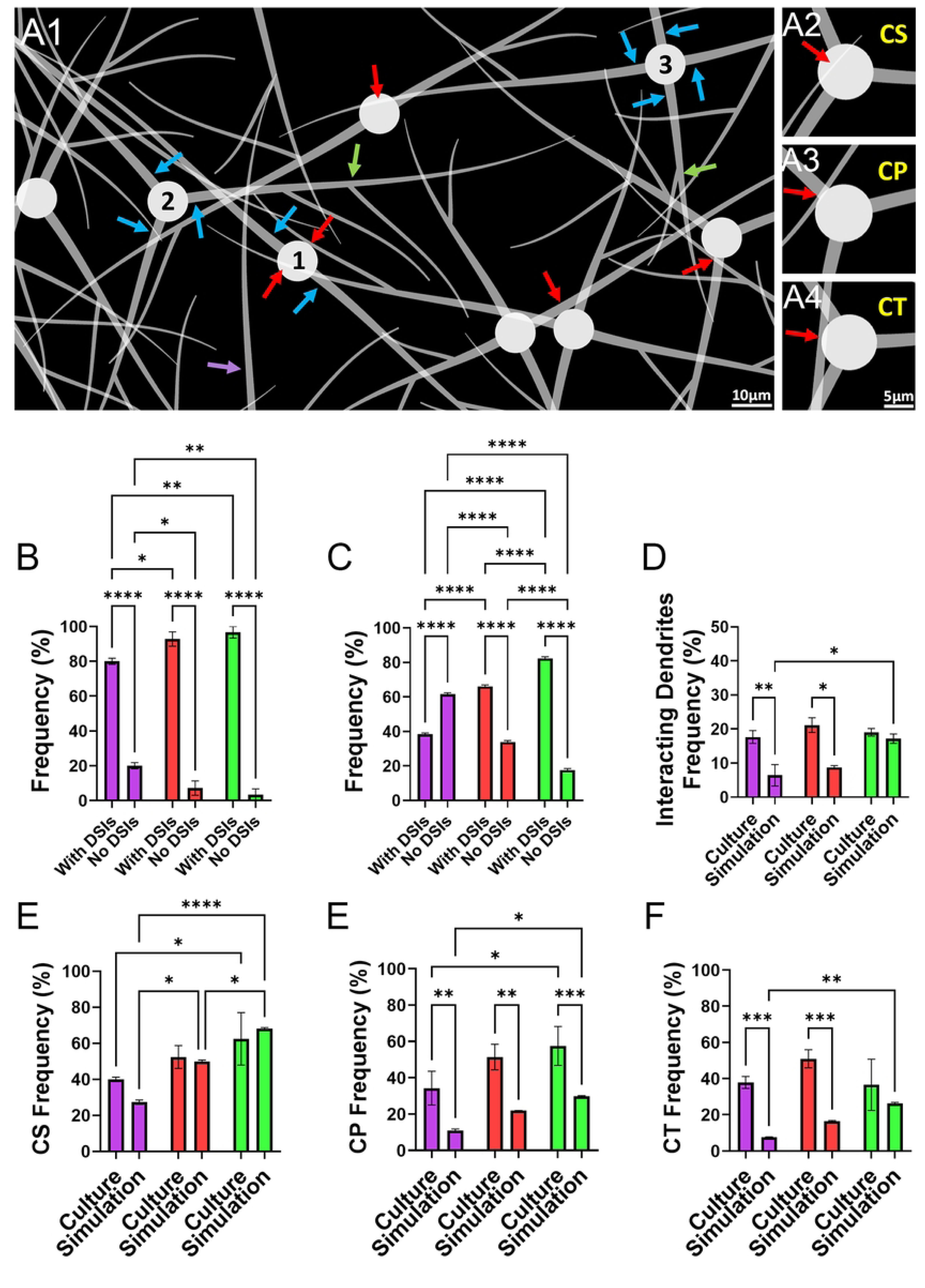
DSI frequencies in hippocampal cultures exceed predictions of a stochastic soma distribution model. (A) Output of the stochastic soma network simulation (SWS-GUI). (A1) Representative simulated field. Three somata are labeled 1–3. Blue arrows indicate the primary dendrites of the labeled cells. Red arrows indicate dendrite–soma interactions (DSIs). Green arrows indicate dendritic branching points. Purple arrow indicates a dendrite belonging to a soma located outside the field. (A2–A4) Magnified examples of the simulated CS, CP, and CT motifs, respectively. (B–G) Comparison of DSI frequencies in cultured networks and in the stochastic simulation. Colors indicate soma density per field: purple, low density (2–6 somata/field); red, medium density (7–9 somata/field); green, high density (10–15 somata/field). (B, C) Soma participation: percentage of somata with at least one DSI versus somata with no DSIs in culture (B) and simulation (C). (D) Dendrite participation: percentage of dendrites forming at least one DSI in culture and simulation. (E–G) Frequencies of the CS (E), CP (F), and CT (G) motifs in culture and simulation. For (D), n represents independent imaging fields (culture: 20, 8, and 3; simulation: 5, 3, and 6 for low-, medium-, and high-density conditions, respectively). For all other panels, data are based on 3 independent replicates, encompassing a total of N = 27–228 observations per condition. Data are presented as mean ± SEM. Statistical significance was determined by two-way ANOVA followed by Tukey’s post hoc test.

To ensure structural realism, the model incorporated somata with a variable number of primary dendrites (range 2–4; Figure 2A1, blue arrows) and utilized randomized parameters to generate branching (green arrows). Furthermore, to account for the complete dendritic population typically observed in imaging fields, the simulation included processes originating from somata positioned outside the defined boundaries (purple arrow). Under these stochastic parameters, the model successfully generated all three morphological DSI subtypes observed in vitro: CS, CP, and CT (Figures 2A2–A4). Notably, these simulated interactions were not restricted to primary dendrites but occurred across different dendritic orders, involving both first- and second-generation branches (Figure 2A1, red arrows). This structural alignment between the stochastic model and biological observations allowed for a direct comparative analysis of DSI frequencies across different network densities.

In hippocampal cultures, the proportion of somas participating in at least one DSI was high even at low cellular densities (2–6 somas per frame), with 80.02% of somas exhibiting DSIs (Figure 2B). This proportion increased with density, reaching 96.66% at densities of 10–15 somas per frame (Figure 2B). In contrast, the stochastic simulation predicted substantially lower interaction frequencies at the above cell densities (38.39% and 66.12%, respectively; Figure 2C). Overall, the observed frequencies in the biological cultures were significantly higher than those predicted by the stochastic model across all densities (P < 0.0001).

To further evaluate DSI frequency, we quantified the percentage of dendrites in each frame that participated in at least one DSI. The mean frequency in culture (averaged across all analyzed fields) remained constant across densities, ranging from 17.66% to 21.15% (Figure 2D). However, the simulation predicted a nearly three-fold lower frequency at 2-6 and 10-15 cell densities (6.42% (p<0.01) and 8.77% (p<0.05), respectively: P <), (Figure 2D).

To determine whether specific DSI subtypes occur more frequently than expected by chance, we compared observed biological frequencies with predictions from a stochastic model across three cellular densities: low (2–6 soma/frame), medium (7–9 soma/frame), and high (10–15 soma/frame). Both CP and CT interactions in culture exhibited substantial enrichment relative to the stochastic model. For CP, frequencies in culture ranged from 34.31% at low density to 57.49% at high density, significantly exceeding simulated values, which ranged from 11.03% to 29.9% across the same densities (Figure 2F). This represents a 2- to 3-fold increase over stochastic predictions (P < 0.01 at low and medium densities; P < 0.001 at high density).

Similarly, CT interactions were more prevalent than predicted by the simulation, with observed values ranging from 37.86% (low density) to 50.95% (high density), compared to predicted ranges of 7.54%–26.28% (Figure 2G). Notably, at the lowest cellular density, the frequency of CT contacts was approximately five-fold greater than the stochastic prediction (P < 0.001 at low and medium densities; P < 0.01 at high density).

By contrast, CS interactions showed no deviation from stochastic predictions. In hippocampal cultures, CS frequencies (ranging from 39.95% at low density to 62.45% at high density) overlapped with the simulated range of 27.29% to 68.1% (P > 0.05; Figure 2E). Statistical comparisons were performed using two-way ANOVA followed by Tukey’s post-hoc test. These results demonstrate that CP and CT interactions occur significantly more frequently in hippocampal cultures than predicted by chance, regardless of cellular density.

### DSI subtype abundancy is developmentally regulated but independent of of neuronal cell adherence or **activity**

Given the significant discrepancies from the random statistical model, we examined the distribution of DSI subtypes CS, CP, and CT across different developmental stages, substrates, and activity levels.

We first quantified the relative frequency of each DSI subtype at 7, 14, and 21 days in vitro (DIV). Frequency was calculated as the percentage of each subtype out of the total number of dendrites forming DSIs (Figure 3A). The frequency of CS interactions significantly decreased as the culture matured, dropping from 39.9% at 7 DIV and 34.56% at 14 DIV to a minimum of 17.39% by 21 DIV (Figure 3A). This final reduction at 21 DIV was significant compared to both 14 DIV (P < 0.05) and 7 DIV (P < 0.01). In contrast, CP interactions showed a significant developmental increase, rising from 34.35% at 7 DIV to 46.91% at 14 DIV and 57.97% at 21 DIV (P < 0.001 for 21 DIV vs. 7 DIV, Figure 3A). The frequency of CT interactions remained stable across all time points, with no significant differences observed between DIVs (Figure 3A).

**Fig 3.**
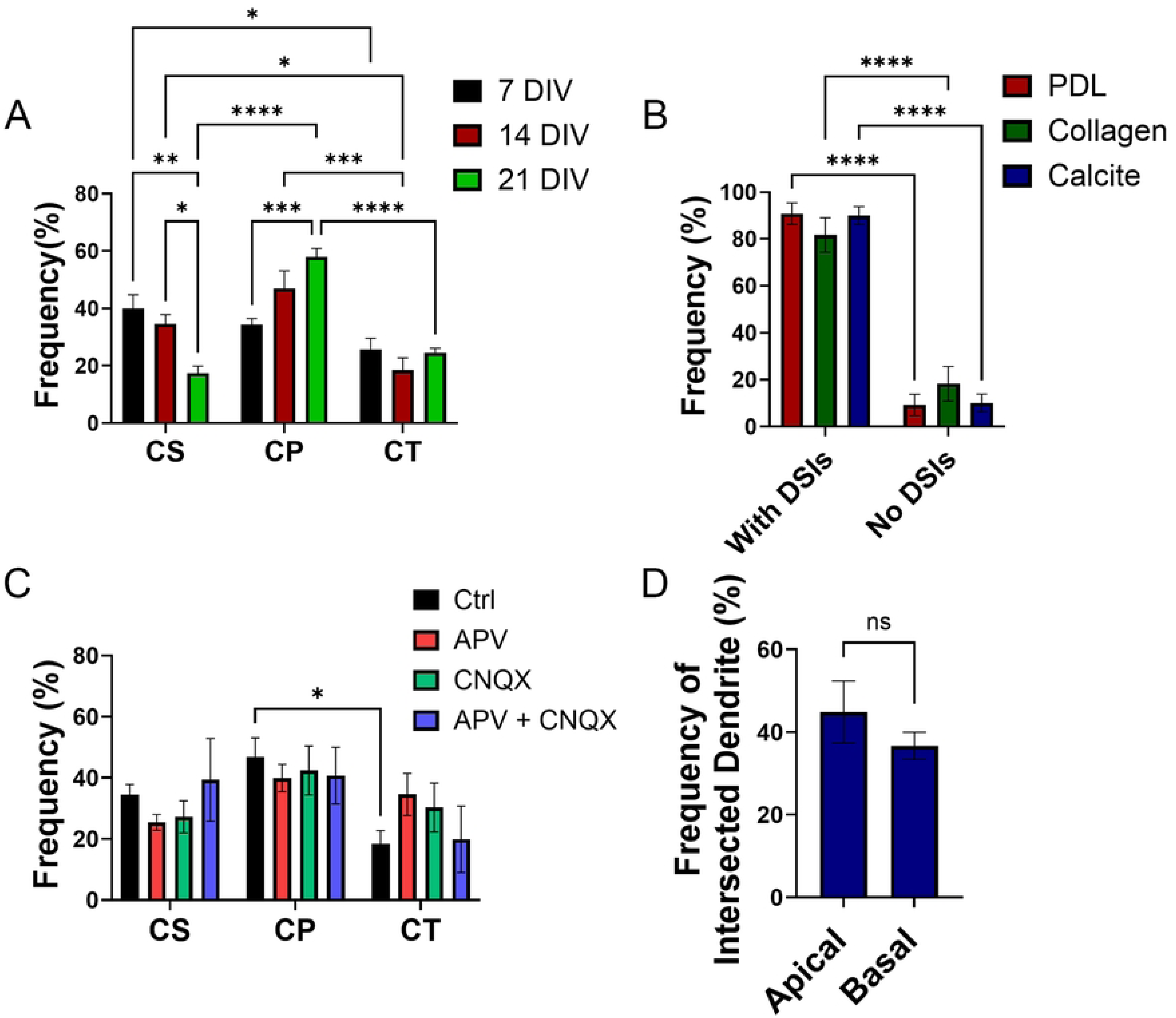
DSI abundancy across developmental stages, substrates, and activity levels. (A) Relative frequency of DSI subtypes at 7, 14, and 21 DIV. (B) Percentage of somas forming DSIs on PDL, Collagen, and Calcite substrates. (C) DSI subtype distribution following pharmacological activity blockade. (D) Frequency of CP interactions on apical versus basal dendritic branches. For all the panels, data reflect n = 3 independent replicates encompassing a total of N = 27–228 observations per condition. Data are presented as mean ± SEM. Statistical significance was determined by one-way or two-way ANOVA followed by Tukey’s post-hoc test.

In addition, at 7 DIV, CS interactions were significantly more frequent than those in CT (P < 0.05). By 14 DIV, significant decreases were observed in CT relative to both CP (P < 0.001) and CS (P < 0.05). Finally, by 21 DIV, CP interactions became significantly more prominent than both CS (P < 0.0001) and CT (P < 0.001, Figure 3A).

To assess whether DSIs formation was influenced by the strength of the cells adhesivity to the culture substrate, we measured the percentage of somas forming at least one DSI on three different substrates: Collagen, Calcite, and Poly-D-Lysine (PDL) (Figure 3B). In all tested substrates, a high percentage of somas participated in DSIs, with frequencies of 81.74% on Collagen 89.95%, on Calcite, and 90.79% on PDL, with no significant differences. For each substrate, the frequency of somas with DSIs was significantly higher than those without DSIs (P < 0.0001, Figure 3B).

We further investigated whether synaptic activity modulates the distribution of DSI subtypes by applying pharmacological blockers (Figure 3C). In this experiment, frequencies were again calculated as the percentage of CS, CP, or CT among the total interacting dendrites. Following treatment with APV, CNQX, or a combination of both, no significant changes were observed in DSI subtype frequencies compared to control cultures (Figure 3C).

To determine whether DSI formation is biased toward a specific dendritic branch type, we analyzed whether CP occurred preferentially on apical vs basal dendritic branches in pyramidal cells (Figure 3D). CP interactions were observed on 44.84% of apical dendrites and 36.66% of basal dendrites. No statistical difference was found between the two dendritic domains.

### DSIs and Dendritic Intersections Display Significantly Higher Density and Intensity of Synaptophysin-Positive Presynaptic Puncta Compared to Non-interacting Segments

Given that synaptic clusters are known to accumulate at dendro-dendritic intersections [16–18]. we were prompted to understand if this holds also for DSI. To achieve that, we first checked if axons are attracted to DSIs (figure 4A1-4A3). Staining with an antibody against NFM revealed no such attraction.

Next, we investigated whether pre-synaptic sites accumulate at the DSIs vicinity. A prominent accumulation of Synaptophysin-positive puncta was detected at the DSI sites (Figure 4B1-4B3), as found also at dendro-dendritic intersections (Figure 4C1-4C3). Quantitative analysis confirmed that both DSIs and dendro-dendritic intersections exhibited significantly higher puncta counts compared to control regions (P < 0.0001; Figure 4D). Notably, no significant difference in puncta density was detected between dendritic intersections and DSIs interactions.

**Fig 4.**
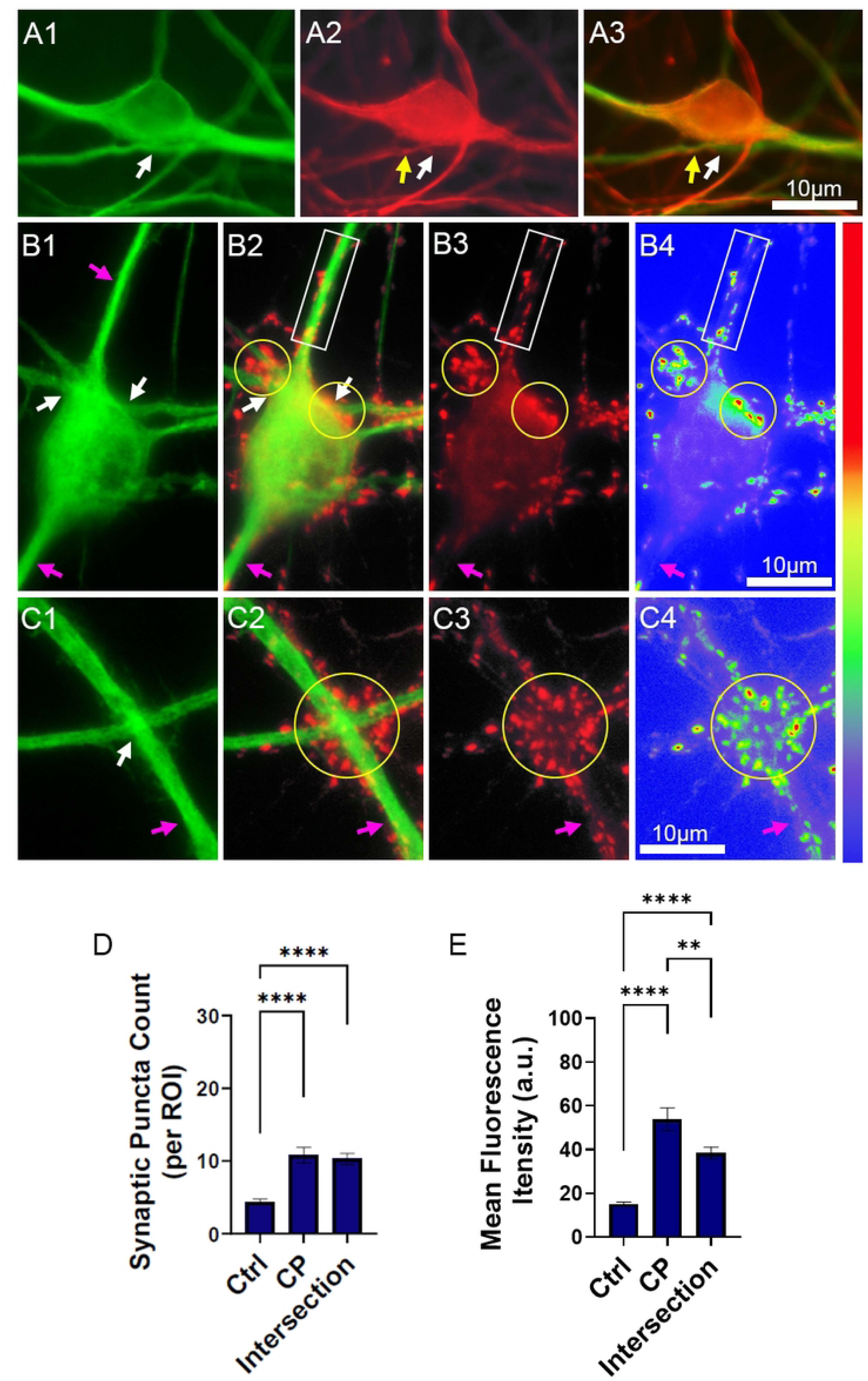
Enhanced density and fluorescence intensity of synaptophysin-positive puncta at DSIs vs. dendro-dendritic intersection sites. (A1-A3) A 7 DIV culture co-stained for MAP2 (green) and NFM (red), with the white arrow indicating to dendrite-soma contacts of a single DSI. (B1-B4) Images showing synaptic clustering near the DSI. All images are of the same field. (B1) A cell body with two contact points pf DSIs (white arrows). Non-intersecting dendritic branches are indicated by purple arrows. (B2) The cell and presynaptic staining (synaptophysin, red). Presynaptic clusters near the DSIs are circled (yellow) and those in non-intersecting dendrites are marked by a white rectangle. (B3) Only synaptophysin staining. (B4) Color-coded image of B3. (C1-C4) Images showing synaptic clustering near a dendro-dendritic intersection. All images are of the same field. (C1) Two intersecting dendritic branches. The point of intersection is indicated by a white arrow. A non-intersecting region is indicated by a purple arrow. (C2) The intersection and presynaptic staining (synaptophysin, red). The presynaptic cluster at the intersection and its vicinity is circled (yellow). (C3) Only synaptophysin staining. (C4) Color-coded image of C3. (D) Quantification of synaptic puncta density. (E) Quantification of mean synaptophysin fluorescence intensity. Sample sizes in D and E: Control (n = 69), CP sites (n = 10), and dendro-dendritic intersections (n = 47). Data represent mean ± SEM. One-way ANOVA with Tukey’s multiple-comparisons test.

We sought to determine if accumulation of presynaptic sites near the DSIs also affected the level of synaptophysin protein in the clustered puncta. A representative color-coded image of the accumulation of Synaptophysin at a DSIs and a dendro-dendritic intersection are shown in Figures 4B4 and 4C4. Quantitative analysis of the mean fluorescence intensity confirmed a significant enrichment of Synaptophysin at both the DSIs and the dendro-dendritic intersections, compared to control (non-DSI non-intersecting dendritic segments (Figure 4E). While the mean intensity in control was 15.2 a.u., the intensity of synaptophysin intensity was significantly higher at both intersecting (CP) sites (53.8 a.u. p < 0.0001) and dendritic intersections (38.7 a.u., p < 0.0001). It was also found that CP sites demonstrated a significantly higher mean fluorescence intensity compared to dendritic intersections (p < 0.01; Figure 4E), suggesting a more pronounced accumulation of presynaptic machinery at dendrite-soma interfaces than at dendrite-dendrite crossovers.

To further examine the mechanisms underlying DSI organization, we next applied minimal quantitative models to the structural and synaptic data. These analyses were designed to test whether the observed subtype distribution of DSIs and the enrichment of synaptic puncta at DSI sites could be explained by stochastic geometry alone or required additional biased formation and local synaptic contributions.

### Minimal mechanistic modeling of DSI structural organization and synaptic enrichment

To examine which minimal processes can account for the structural organization of DSIs, we modeled the observed frequency of each DSI subtype as:

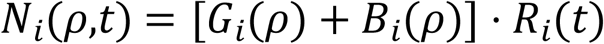

where 𝑁_𝑖_(𝜌,𝑡) is the observed frequency of subtype 𝑖 at cellular density 𝜌 and developmental time 𝑡, 𝐺_𝑖_(𝜌) is the stochastic expectation derived from the random spatial simulation, 𝐵_𝑖_(𝜌) is a biased-formation term representing directed growth beyond stochastic expectation, and 𝑅_𝑖_(𝑡) is a developmental refinement term. Under this framework, the stochastic model corresponds to 𝐵_𝑖_ = 0 and 𝑅_𝑖_ = 1; the stabilization-only model corresponds to 𝐵_𝑖_ = 0; the directed-growth-only model corresponds to 𝑅_𝑖_ = 1; and the combined model includes both terms. This formulation was used to compare the relative ability of formation bias and stabilization to account for the observed subtype distribution of CS, CP, and CT.

At low cellular density (2–6 soma/frame), the directed-growth-only model reproduced the observed enrichment of CP and CT more closely than the stabilization-only model, whereas CS remained less well explained by directed growth alone (Figure 5A). At high cellular density (10–15 soma/frame), the same pattern was observed: the directed-growth-only model again approximated the observed frequencies of CP and CT better than the stabilization-only model, while CS remained closer to the stochastic baseline (Figure 5B). In contrast, the stabilization-only model consistently overestimated subtype frequencies, particularly for CP and CT at high density. These comparisons indicate that biased formation provides a better minimal explanation for the structural enrichment of CP and CT than stabilization alone. This interpretation is consistent with the experimental findings that CP and CT exceed stochastic expectation, whereas CS largely overlaps with the random prediction, and with the developmental shift in which CP increases over time, CS decreases, and CT remains relatively stable.

**Figure 5.**
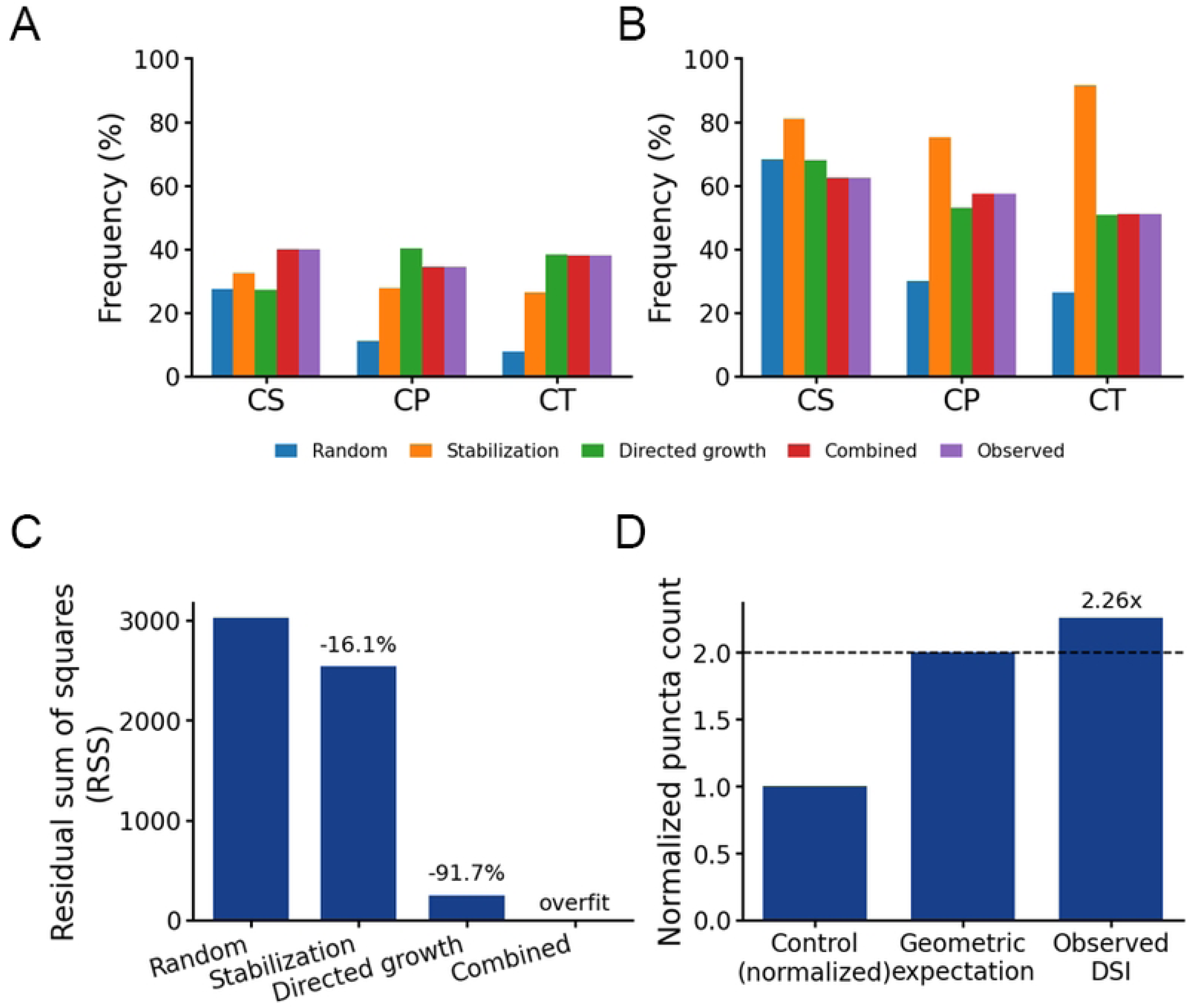
Quantitative models of DSI organization and synaptic puncta enrichment. (A) Comparison of model predictions and observed frequencies of DSI subtypes at low cellular density (2–6 soma/frame). (B) Same analysis as in (A), shown for high cellular density (10–15 soma/frame). Panels A and B share a common legend. (C) Comparison of model performance across density conditions, expressed as RSS. Values above the bars indicate the reduction in residual error relative to the stochastic baseline. The combined model reproduces the observed values exactly because it contains sufficient flexibility to fit all data points. (D) Normalized synaptic puncta count at DSI sites relative to control dendritic segments. Control was normalized to 1, and the expected value for simple geometric superposition was set to 2. The observed puncta count at DSI sites was higher than control and slightly above the geometric expectation.

To compare model performance quantitatively, we calculated the residual sum of squares (RSS) for each model across density conditions. Relative to the stochastic baseline, the stabilization-only model reduced the overall residual error by 16.1%, whereas the directed-growth-only model reduced it by 91.7% (Figure 5C). The combined model reproduced the observed values exactly, but because it contains sufficient flexibility to fit all data points directly, it should be interpreted as an overfit reference rather than as a uniquely supported mechanistic explanation. Together, these results support the interpretation that early DSI organization is shaped predominantly by biased formation, whereas later developmental changes are consistent with subtype-specific refinement rather than with stabilization alone.

To evaluate whether synaptic puncta enrichment at DSI sites could be explained by geometric overlap alone, we modeled puncta abundance at DSI sites as:

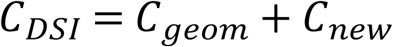

where 𝐶_𝐷𝑆𝐼_ is the puncta count at a DSI site, 𝐶_𝑔𝑒𝑜𝑚_ is the puncta count expected from simple geometric superposition of the interacting compartments, and 𝐶_𝑛𝑒𝑤_ is an additional local contribution. Under the simplifying assumption that the interacting compartments contribute similar baseline puncta density, the geometric expectation can be approximated as:

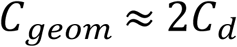

where 𝐶_𝑑_ is the puncta count at a control dendritic segment. We therefore defined the normalized enrichment ratio as:

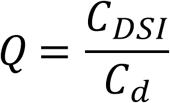

for which 𝑄 = 2 represents the value expected from simple geometric superposition. Using the puncta-count quantification, the observed enrichment at DSI sites was 2.26-fold relative to control (Figure 5D), slightly exceeding the geometric expectation. This result suggests that puncta enrichment at DSI sites is not fully accounted for by passive spatial overlap alone and may include an additional local synaptic contribution.

Overall, these results show that DSIs are common and structurally diverse features of cultured hippocampal neurons. CP and CT interactions occurred more frequently than expected from stochastic simulations, whereas CS frequencies were largely consistent with random predictions, and the relative abundance of the subtypes changed across development. CP sites were also associated with increased synaptophysin puncta density and intensity, and minimal quantitative modeling indicated that the observed DSI organization is better explained by biased formation with developmental refinement than by stochastic organization alone.

## Discussion

The present study identifies DSIs as a non-random class of heteroneuronal structural motifs in hippocampal cultures. DSIs occur at frequencies that exceed stochastic predictions and display subtype-specific organization, with CP and CT selectively enriched relative to CS. In addition, DSI composition is developmentally dynamic, with CP progressively emerging as the dominant configuration over time. This combination of non-random occurrence and subtype-specific refinement is consistent with growing evidence that local neuronal contacts are not distributed uniformly, but are shaped by structural and developmental constraints that bias connectivity patterns [5].

Functionally, this structural bias is accompanied by increased presynaptic protein accumulation at DSI sites, indicating that these configurations are not only overrepresented but also preferentially associated with synaptic organization [22,23]. Similar coupling between local geometry and synaptic clustering has been described in other systems, where spatially defined contact sites serve as preferred loci for presynaptic assembly and stabilization [24,25]. Within this context, the present findings suggest that DSIs - and in particular CP - represent structurally defined sites that bias the localization of presynaptic components.

Together, these results support the interpretation that DSI organization cannot be explained by stochastic geometry alone. Consistent with the minimal modeling in Fig. 5, the enrichment of CP and CT is better accounted for by biased formation than by stabilization alone, whereas CS remains closer to the stochastic baseline. The same analysis further suggests that later developmental changes in subtype composition reflect subtype-specific refinement, and that synaptic puncta enrichment at DSI sites is not fully explained by geometric overlap alone.

### 1. DSIs represent non-random, geometry-dependent structural motifs within a broader class of contact-mediated organization

A central result of the present study is that DSIs occur at frequencies that exceed those predicted by a stochastic geometric model, indicating that their formation cannot be explained by random spatial overlap alone. Deviations of this kind have been widely reported in neuronal systems, where contact formation reflects an interplay between spatial constraints and regulated developmental processes rather than purely incidental encounters [5,26,27]. These findings position DSIs within a broader framework in which local neuronal contacts are biased beyond stochastic expectations.

The density-dependent behavior of DSIs further reinforces this interpretation. At low cellular densities, where stochastic models predict relatively few interactions, the discrepancy between observed and expected frequencies is maximal. As density increases, incidental contacts become more probable, partially masking this deviation. This relationship suggests that biological contributions to contact formation are most clearly revealed under sparse conditions, whereas at higher densities, geometric constraints increasingly dominate. Similar interactions between structural constraints and connectivity rules have been described in dense neuronal reconstructions, where wiring patterns emerge from the combined effects of spatial proximity and selective stabilization [28,29].

Within this non-random regime, DSIs also exhibit subtype-specific organization. CP and CT configurations are enriched relative to stochastic expectation, whereas CS interactions largely conform to random predictions. This selective enrichment indicates that not all geometrical configurations are equally favored, but rather that specific spatial arrangements are preferentially formed and subsequently refined. Comparable selectivity has been observed in analyses of dendritic organization, where only particular configurations of neighboring branches give rise to stable interaction sites [18,30,31].

This interpretation is reinforced by the minimal model comparison in Fig. 5. At both low and high cellular densities, the directed-growth-only model reproduced the observed enrichment of CP and CT more closely than the stabilization-only model, whereas CS remained closer to the stochastic prediction. Consistent with this pattern, the directed-growth-only model reduced the residual error by 91.7% relative to the stochastic baseline, compared with 16.1% for the stabilization-only model. Although the combined model fit the data exactly, its flexibility suggests that it should be interpreted as an overfit reference rather than as a uniquely supported mechanism. Together, these comparisons support biased formation as the main contributor to early DSI organization.

Importantly, similar non-random enrichment has been described for interactions between dendritic branches, where neighboring branches regulate their spatial proximity and converge to form clustered contact regions. These regions exhibit elevated densities of contacts and synaptic components and occur more frequently than predicted by random models, indicating regulated rather than incidental formation [16–18]. Furthermore, such interactions are influenced by neuronal activity and are associated with local strengthening of connectivity, suggesting that dendritic proximity itself serves as a structural mechanism for organizing synaptic distribution and network efficiency [16–18]. The parallels with the present findings suggest that DSIs extend this general principle of geometry-dependent, non-random contact formation to dendrite–soma interfaces.

### 2. Developmental refinement selectively favors CP as a structurally privileged configuration

DSI composition is not static, but undergoes a marked developmental shift characterized by a progressive increase in CP alongside a reduction in CS, while CT remains relatively stable. This divergence indicates that subtype distribution is not determined solely at the stage of initial contact formation but is actively refined over time. This interpretation is also consistent with the quantitative analysis in Fig. 5, which suggests that the early enrichment of CP and CT is better explained by biased formation, whereas the later divergence between CP and CS is more consistent with subtype-specific developmental refinement than with stabilization alone.

Developmental refinement of neuronal contact is a well-established feature of circuit maturation, in which early structural interactions are selectively stabilized and refined rather than uniformly preserved [32]. In the present study, and consistent with the minimal analysis in Fig. 5, the increase in CP together with the decline in CS is more consistent with subtype-specific refinement of initially biased contacts than with stabilization alone.

A key structural distinction of CP is that it is the only configuration that involves direct contact between dendritic branches. This feature aligns it with previously described interactions between neighboring dendrites, which were shown to form stable, non-random contact sites and to reflect regulated spatial organization rather than incidental proximity [16–18]. In addition, CP is typically located near proximal dendritic regions, a spatial domain known to play a central role in shaping dendritic interactions and structural organization [33,34]. This positional bias may further contribute to the preferential enrichment and maintenance of CP over other configurations.

Taken together, these observations suggest that developmental processes bias DSI organization toward configurations that incorporate an interaction composed of dendrite-soma and dendro-dendritic contacts, thereby selectively enriching CP within the DSI population.

### 3. DSIs are focal sites of synaptic clustering and strengthening

DSIs are consistently associated with regions of increased synaptic density, forming localized clusters of synapses whose density and strength are higher than those observed in surrounding dendritic regions. This spatial coupling between structural contact sites and synaptic accumulation indicates that DSIs represent preferential loci for synaptic organization within the network. Synaptic clustering is a well-established organizational principle in neuronal systems, with significant implications for signal integration [9,10]. When synapses are spatially grouped, the reduced distance between them enhances local cooperativity and increases the probability of temporal and spatial summation. Such clustered arrangements promote nonlinear integration within dendritic segments and enable coordinated activation of neighboring inputs [9,10]. In addition, clustered synapses have been associated with increased synchrony of activation and the emergence of burst-like firing patterns, reflecting the concerted recruitment of nearby synaptic inputs [9,10]. These properties indicate that synaptic clustering is not merely a structural feature, but a key determinant of neuronal computation.

This interpretation is further supported by the minimal analysis in Fig. 5D. When puncta abundance at DSI sites was normalized to control dendritic segments, the observed enrichment reached 2.26-fold, slightly above the two-fold value expected from simple geometric superposition. This result suggests that synaptic puncta accumulation at DSI sites is not fully explained by passive geometric overlap and is consistent with an additional local contribution to presynaptic organization.

Beyond their spatial organization, DSI-associated sites also exhibit increased synaptophysin intensity, indicating enhanced presynaptic enrichment. The relationship between spatial clustering and local synaptic strengthening has been extensively documented, with nearby synapses often undergoing coordinated potentiation and exhibiting enhanced transmission efficacy [4,9,10]. In the present study, however, the relevant experimental measure was presynaptic protein accumulation rather than direct physiological strength, and the increased synaptophysin signal is therefore more conservatively interpreted as evidence of enhanced presynaptic specialization at DSI sites.

Similar relationships between local structural configurations and synaptic enrichment have been described in interactions between neighboring dendritic branches, where contact sites are associated with elevated synaptic density and increased functional output. In those systems, regions of dendritic proximity exhibit both higher synaptic density and enhanced synaptic efficacy, closely paralleling the properties observed at DSIs [16–18]. Moreover, the quantitative comparisons presented in the current study indicate that the magnitude of synaptic enrichment at DSIs is comparable to that observed at dendritic intersection sites, reinforcing the idea that both structures share common organizational principles. These parallels suggest that DSIs and dendrite–dendrite interaction sites represent related structural motifs that support clustered and potentiated synaptic organization within neuronal networks.

A key observation in the present study is the absence of corresponding axonal enrichment at DSIs, despite the clear increase in synaptic density and presynaptic enrichment. This dissociation suggests that synaptic accumulation at these sites is not driven by preferential convergence of axons, but rather reflects local processes that promote synapse formation or stabilization. Previous work has shown that synaptogenesis can be regulated by local structural cues and contact-dependent mechanisms, independently of large-scale axonal targeting [35,36]. Within this framework, DSIs may provide a structural environment that facilitates the local generation and strengthening of synaptic connections.

### 4. Implications and conceptual framework

The findings of the present study support a model in which DSIs represent structurally defined sites that contribute to the spatial organization of synaptic connectivity. By combining non-random formation, developmental refinement, and association with synaptic clustering and strengthening, DSIs emerge as elements that link local geometry with functional network properties.

Within this framework, DSIs may serve as organizational units that locally bias the distribution and efficacy of synaptic inputs. Rather than reflecting incidental contacts, these structures appear to define regions in which synaptic organization is both spatially constrained and functionally enhanced. This interpretation is consistent with broader principles of neuronal organization, in which connectivity is shaped by an interplay between structural constraints and activity-dependent processes [4,5].

Importantly, the parallels observed between DSIs and interactions between dendritic branches suggest that common geometrical principles underlie multiple forms of local connectivity. The extension of these principles to dendrite - soma interfaces indicates that such organization is not limited to a specific class of interactions, but may represent a general mechanism for structuring synaptic networks.

The structural and functional features described here suggest that DSIs provide a substrate for localized modulation of synaptic organization. Future studies will be required to determine the extent to which these structures contribute to network-level dynamics and plasticity under physiological conditions.

### 5. DSI formation is independent of synaptic activity and differs from activity-dependent dendritic interactions

An additional key finding of the present study is that DSI formation is not significantly affected by variations in substrate conditions or by pharmacological inhibition of synaptic activity. The persistence of DSIs under these conditions indicates that their formation does not depend on specific extracellular substrates or ongoing synaptic transmission.

The lack of sensitivity to substrate suggests that DSI formation is not driven by externally imposed adhesion differences or surface-dependent growth constraints. Instead, it supports a model in which DSIs emerge from intrinsic structural interactions between neurons, consistent with studies showing that neurite growth and organization are governed by cell-intrinsic mechanical and cytoskeletal properties rather than solely by external substrate cues [13,37]. Notably, no significant differences were observed in DSI frequency between apical and basal dendritic compartments of pyramidal neurons, further indicating that DSI formation is not restricted to specific dendritic domains.

Equally important is the observation that blocking synaptic activity does not alter DSI formation. This distinguishes DSIs from other forms of dendritic organization, including interactions between neighboring dendritic branches, which have been shown to be at least partially regulated by neuronal activity [16–18]. Activity-dependent mechanisms have been implicated in the stabilization, refinement, and spatial organization of dendritic contacts and synaptic clustering [5,38]. In contrast, the activity-independence of DSIs suggests that they arise through mechanisms that precede or operate independently of synaptic transmission.

Together, these findings indicate that DSIs represent a structural framework that is established independently of both external substrate conditions and neuronal activity, and may therefore provide a pre-existing scaffold upon which activity-dependent processes subsequently act.

### 6. Limitations and implications

Several limitations of the present study should be considered when interpreting the findings. First, the analysis was conducted in dissociated hippocampal cultures, which, while allowing precise structural and functional characterization, do not fully recapitulate the spatial organization and connectivity constraints present in intact brain tissue. As such, the extent to which DSIs form and operate within in vivo circuits remains to be determined.

Second, although DSIs are associated with increased synaptic density and presynaptic enrichment, the present study does not directly address the molecular mechanisms underlying their formation or maintenance. In particular, it remains unclear which adhesion molecules or signaling pathways mediate the establishment of these structures, and whether distinct molecular programs differentiate DSIs from other forms of neuronal contact.

In addition, while the functional consequences of synaptic clustering near DSIs are inferred based on established principles of synaptic integration and plasticity, direct measurements of how DSIs influence neuronal output or network activity were not performed. Future studies combining structural analysis with functional recordings will be required to determine the extent to which DSIs modulate information processing at the cellular and network levels.

Despite these limitations, the present findings have several important implications. The identification of DSIs as non-random, structurally defined sites associated with synaptic clustering and presynaptic enrichment suggests that local neuronal geometry plays a central role in shaping synaptic organization. The independence of DSI formation from substrate conditions, synaptic activity, and dendritic domain further supports the idea that these structures arise from intrinsic organizational principles rather than externally imposed constraints.

Together, these observations position DSIs as candidate structural units that contribute to the spatial patterning and functional modulation of synaptic networks. By linking dendritic organization, contact geometry, and synaptic clustering, DSIs may represent a general mechanism through which neuronal circuits achieve localized plasticity and efficient information integration.

## S1_Supporting_Information

Compendium of raw data and statistical summaries.

This ZIP file contains individual Excel workbooks with raw data for each figure, a summary of sample sizes, and a detailed guide to the data organization.

